# A probabilistic framework for decoding behavior from in vivo calcium imaging data

**DOI:** 10.1101/827030

**Authors:** Guillaume Etter, Frederic Manseau, Sylvain Williams

## Abstract

Understanding the role of neuronal activity in cognition and behavior is a key question in neuroscience. Previously, *in vivo* studies have typically inferred behavior from electrophysiological data using probabilistic approaches including Bayesian decoding. While providing useful information on the role of neuronal subcircuits, electrophysiological approaches are often limited in the maximum number of recorded neurons as well as their ability to reliably identify neurons over time. This can be particularly problematic when trying to decode behaviors that rely on large neuronal assemblies or rely on temporal mechanisms, such as a learning task over the course of several days. Calcium imaging of genetically encoded calcium indicators has overcome these two issues. Unfortunately, because calcium transients only indirectly reflect spiking activity and calcium imaging is often performed at lower sampling frequencies, this approach suffers from uncertainty in exact spike timing and thus activity frequency, making rate-based decoding approaches used in electrophysiological recordings difficult to apply to calcium imaging data. Here we describe a probabilistic framework that can be used to robustly infer behavior from calcium imaging recordings and relies on a simplified implementation of a naive Baysian classifier. Our method discriminates between periods of activity and periods of inactivity to compute probability density functions (likelihood and posterior), significance and confidence interval, as well as mutual information. We next devise a simple method to decode behavior using these probability density functions and propose metrics to quantify decoding accuracy. Finally, we show that neuronal activity can be predicted from behavior, and that the accuracy of such reconstructions can guide the understanding of relationships that may exist between behavioral states and neuronal activity.

## Introduction

Early *in vivo* studies have established relationships between external variables and neuronal activity, including (but not restricted to) auditory information in the auditory cortex (Katsuki et al., 1956), visual stimuli in the visual cortex (Hubel and Wiesel, 1962), and spatial information in the hippocampus (O’Keefe and Dostrovsky, 1971). Based on the widely influential information theory (Shannon, 1948), it has previously been proposed that neurons can act as ‘communication channels’ between physiological variables (input) and spike trains (output) (Richmond and Optican, 1990; Richmond et al., 1990; Skaggs et al., 1993). In addition to providing metrics to quantify the extent to which external variables can be encoded in neurons, these studies laid the first foundations in establishing computational tools to predict animal behavior merely using neuronal activity. This process, termed decoding, is essential in understanding the role of neuronal activity in behavior, and the success rate of predictions can be used as a metric of understanding of a given system. Among techniques that have been described in this context, Bayesian decoding in particular has been relatively popular and widely used (Brown et al., 1998; Zhang et al., 1998; Gerwinn, 2009; Quian Quiroga and Panzeri, 2009; Koyama et al., 2010).

While the literature on *in vivo* neuronal physiology has been largely dominated by electrophysiological studies, calcium imaging methods have recently gained popularity. Originally performed at the single cell level with the aid of calcium sensors (Grynkiewicz et al., 1985; Persechini et al., 1997), calcium imaging can now be performed *in vivo*, in large neuronal assemblies, and over very long periods of time (Ziv et al., 2013; Sheintuch et al., 2017; Gonzalez et al., 2019). These major improvements coincided with the development of genetically encoded calcium indicators (GECI), including the popular GCaMP (Nakai et al., 2001; Tian et al., 2009; Ohkura et al., 2012; Chen et al., 2013). In recent years, calcium imaging methods have seen the development of various computational tools that solve the problem of signal extraction from raw calcium imaging video recordings. In particular, several groups have proposed open-source software codes to perform fast, recursive motion correction (Pnevmatikakis and Giovannucci, 2017), offline (Pnevmatikakis et al., 2016; Zhou et al., 2018) and online (Giovannucci et al., 2017) extraction of neuronal spatial footprints and their associated calcium activity, temporal registration of neurons across days (Sheintuch et al., 2017), and complete analysis pipelines have been made available (Giovannucci et al., 2018). The aforementioned open source codes have significantly facilitated the analysis of calcium imaging datasets. Most often, one of the objectives when using such a tool is to understand the neural basis of behavior. Unfortunately, there are only a few open source analysis toolboxes that can relate calcium imaging data to behavior to this day (Tegtmeier et al., 2018; www.miniscope.org). While these useful analytical tools allow the exploration of relationships between calcium signals and behavior, they are mostly restricted to visualization and correlation. Previous studies have made use of naive Bayesian classifiers to infer rodent behavior from calcium imaging data recorded in the motor cortex (Huber et al., 2012; Kondo et al., 2018), hippocampus (Ziv et al., 2013; Mau et al., 2018; Gonzalez et al., 2019), or amygdala (Grewe et al., 2017). While these analyses usually result in accurate predictions of behaviors, there is no consensus on the methodology, and in particular the input signal to the classifier, or other preprocessing steps (e.g. smoothing of neuronal tuning curves used by the classifier).

While calcium imaging does not allow the determination of exact spike timing, some methods have been proposed to approximate spiking activity from calcium imaging data by deconvolving calcium transients (Deneux et al., 2016; Pachitariu et al., 2018; Rahmati et al., 2018). Consequently, one strategy that can be employed is to first estimate deconvolution parameters from ground truth data (e.g. *in vitro* unit recording in brain slices combined with calcium imaging) to then apply them to recordings performed *in vivo*. However, one major caveat with this approach is that physiological responses can differ greatly between *in vivo* and *in vitro* conditions (Belle et al., 2018) leading to erroneous parameter estimation. Another obstacle to using deconvolved signals and estimated spikes to decode calcium activity is that the very nature of calcium imaging does not allow to determine exact spike timing. While unit recordings are typically done at sampling rates exceeding 10 KHz, 1-photon microendoscopes used in freely moving animals usually sample images at 30 frames per second or less, and spike trains will generally be associated with large calcium transients of varying size and duration. Consequently, one could for example correctly estimate that a neuron fires 10 action potentials based on the observation of a single calcium transient, however the exact timing of each spike would remain unknown, and could happen anywhere within a ~33 ms window (for calcium imaging performed at 30 Hz).

Importantly, another issue encountered when performing calcium imaging with GCaMP is photobleaching, which leads to a progressive loss of signal due to the destruction of fluorescent proteins that report calcium influx. Unlike electrophysiological unit recordings that can be performed for several hours, calcium imaging is thus typically performed for shorter durations. While it is possible to follow GCaMP-positive cell assemblies over months (Ziv et al., 2013; Sheintuch et al., 2017), each recording session has to be limited in duration to avoid photobleaching. This results in low sampling that can be problematic when trying to associate neuronal activity with a certain behavior: some behavioral states can be over- or underrepresented and concurrently, calcium activity can be too sparse to establish tuning curves of neuronal activity.

Here we propose simple analytical methods to relate calcium activity to behavior by (1) extracting periods of activity in calcium imaging data without approximating spike timing and subjecting actual data to null hypothesis testing in order to solve the problem of low sampling, (2) decoding behavior by using previously computed probability density functions in a naive Bayesian classifier, and (3) reconstructing neuronal activity from behavior and assessing the quality of neuronal coding.

## Results

### 1. Establishment of probabilistic neural tuning curves

To demonstrate the usefulness of our method, we performed calcium imaging in a well characterized system: CA1 pyramidal cells of the dorsal hippocampus (fig. 1a). These neurons are known to display spatial tuning and are commonly referred to as place cells (O’Keefe and Dostrovsky, 1971). We trained a mouse to run back and forth on a 100 cm long linear track by providing sucrose water rewards at each end of the track and scheduling homecage access to water every day (fig. 1b). We recorded ~400 neurons in these conditions (fig. 1c). After extracting neuronal spatial footprints (fig. 1d), we visualized corresponding calcium activity along with the position and locomotor speed of the animal (fig. 1e). Previous studies have shown that immobility periods are associated with replay of experience (Foster and Wilson, 2006; Diba and Buzsáki, 2007; Davidson et al., 2009). In order to focus on the spatial tuning curves of CA1 neurons, we therefore excluded periods of immobility (< 5cm.s^-1^) that could potentially contain periods of neuronal activity that reflect tuning to internal, rather than external variables.

**Figure 1.**
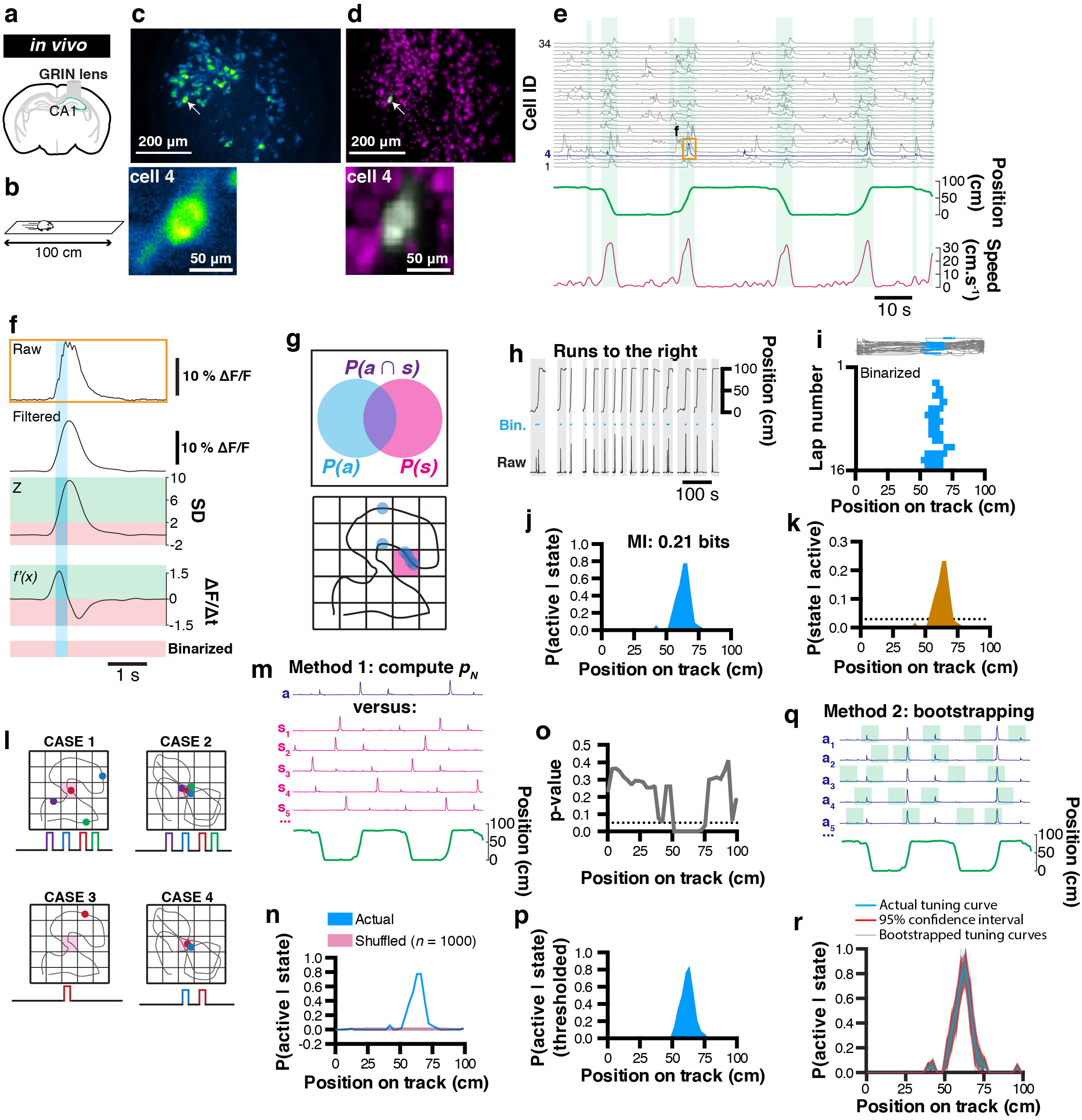
Rationale for extracting spatial coding characteristics of CA1 pyramidale cells. a, diagram of GRIN lens implanted over CA1 pyramidal cells of the dorsal hippocampus. b, calcium imaging was performed as a mouse was running in alternating directions on a linear track. c, maximum projection of the corresponding field of view. d, corresponding extracted spatial footprints using CNMFe. e, example traces from a subset of extracted cells aligned with position on a linear track and locomotor speed. Running epochs are indicated with green stripes. f, example raw transient (top) from one cell and corresponding filtered, z-scored, first-derivative, and binarized signals. g, rationale used to extract unconditional and joint probabilities from binarized data that can be later used to compute conditional probabilities. h, mouse location on the linear track with corresponding raw calcium activity and derived binary trace (blue). Only runs to the right are considered here. i, (top) mouse trajectory on the linear track (gray) with corresponding locations where one cell’s binarized activity was detected (blue dots), and (bottom) location of binarized activity on the linear track for each run (n = 16 runs). j, probability P(active|state) of cell #4 to be active given the location on the linear track, and corresponding mutual information (MI, top). k, derived posterior probability of the mouse being in a location given cell activity P(state|active) (ocher) compared to uniformity (dotted line). l, example cases of poor variable coding (case 1), superior variable coding (case 2), poor variable coding with sparse information (case 3), and superior variable coding with sparse information (case 4). m, method for computing a p-value: actual (a) calcium trace, corresponding circular permutations (s_n_), and corresponding location (green). n, probability P(active|state) of cell #4 being active given location (blue) and corresponding average shuffled distribution from n = 1000 surrogates (the thickness of the line represents the SEM). o, p-value computed using actual data and shuffled surrogates for each location of the linear track. p, thresholded place field retaining only significant P(active|state) values. q, method for computing 95% confidence interval from bootstrap samples. r, actual tuning curve P(active|state) (blue) and corresponding 95% confidence interval (red) computed from bootstrap samples (gray; n = 1000).

In order to compute probabilities that will be used in later analyses of tuning curves, we sought to discriminate periods of activity versus inactivity. To this end, we devised a simple binarizing method where raw calcium signals are first filtered (to remove high frequency fluctuations that could erroneously be detected as transient rise periods), and we considered periods of activity as following the two criteria: (1) the signal amplitude of a normalized trace has to be above 2 standard-deviations, and (2) the first order derivative has to be positive (thus corresponding to a transient rise period; fig. 1f).

#### 1.1 Extracting probability values in a Bayesian context

Following the binarization of raw calcium traces, we propose a probabilistic framework to describe how the activity of a neuron encodes a certain behavior or state (fig. 1g). To this end, we can first compute the probability of a neuron to be active *P(A)* using the following formula:

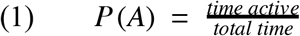

*P(A)* only informs on the activity rate of a neuron over the course of a recording session and corresponds to the marginal likelihood in a Bayesian context. We can also compute the probability of spending time in a given behavioral state *i*:

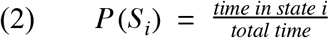

*P(S)* can be informative on whether the distribution of behavioral states is homogeneous or inhomogeneous, which can potentially lead to biases in further analyzes. In a Bayesian context, *P(S)* corresponds to the prior. Additionally, the joint probability *P*(*S_i_⋂ A*) of a given neuron to be both active and in a given state *i* will be later used to compute information metrics:

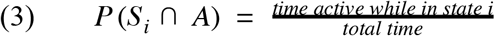

We can then compute the probability that a cell is active given the animal is in a state *i*:

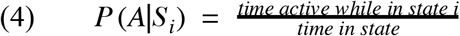

*P* (*A*|*S*) is more informative and can be interpreted as a tuning curve. In a Bayesian framework, this probability is also known as the likelihood. For example, a probability value of 0.8 means that a cell is active 80 % of the time when the animal is in a given behavioral state.

In our example, we isolated running periods when the mouse was running towards the right hand side of the linear track (fig. 1h), and divided the track in 3 cm bins. Each bin thus represents a discrete state, and while visualizing the activity of neuron #4 for each run, it is apparent that this cell displays some spatial tuning (fig. 1i). We thus computed *P* (*A*|*S*) for that cell and found a peak likelihood of 0.78 at ~64.5 cm from the left hand side of the track (fig. 1j). Finally, using classical Bayesian inference, we can infer the probability that the animal is in a state *S_i_* given neuronal activity A:

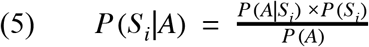

*P* (*S*|*A*) is the posterior probability distribution of states given neuronal activity and will be used later on to decode behavior.

### 1.2 Testing significance of tuning curves

One current issue with calcium imaging is photobleaching, which prevents extended recordings and thus restricts the sampling of both neuronal activity and behavioral data. Experimenters can thus be frequently faced with one of four cases: first, sampling of both behavior and neuronal activity are sufficient, and there is no apparent relationship between these two variables (fig. 1l, case 1). Secondly, sampling is sufficient and there is a very clear relationship between behavior and neuronal activity (fig. 1l, case 2). Thirdly, sampling is too low to observe a clear phenomenon (not enough coverage of behavioral states, sparse neuronal activity; fig. 1l, case 3). Lastly, behavioral sampling is sufficient, but neuronal activity is sparse and while there is an apparent relationship between behavior and neuronal activity, there is no absolute confidence that this could indeed be the case (fig. 1l, case 4). In every case, we will want to test whether the tuning curves that have been derived are significantly different from chance.

#### 1.2.1 Deriving p-values from shuffled distributions

One solution we propose to this problem is to confront the actual data to a null hypothesis that there is no relationship between behavior and neuronal activity. To this end, we generated a distribution of tuning curves that are computed from the actual calcium trace, but that has been shifted in time so that any potential relationship to behavior is lost. Circular permutations can be used to remove the temporal relationship that exists between neuronal activity and behavior; fig. 1m). We recommend this shuffling method because it preserves the temporal structure of calcium transients and leads to more conservative results, as opposed to a complete randomization of every data point which often gives rise to non-physiological data, and thus inflates the significance value of results. The choice of a null hypothesis should however be determined carefully depending on the nature of the question asked. In our example, we performed *n* = 1000 random circular permutations, computed the mean as well as standard error of the mean (SEM), and compared it to our actual tuning curve (fig. 1n). Because the shuffled data points cannot be assumed to be distributed normally, transforming the actual data into a z-score cannot be used to derive a measure of significance. Instead, one can compute a p-value that corresponds to the number of data points from the shuffled distribution that are greater than the actual data, divided by the number of permutations (Cohen, 2014). Note that using this method, the p-value can take null values in the event where all data points from the shuffled distribution fall under the actual data. After computing a p-value value from our actual data and shuffled distribution (fig. 1o), a threshold (typically of 0.05 or 0.01) can be used to display only significant data points (we used a 0.05 threshold in our example; fig. 1p).

#### 1.2.2 Deriving confidence intervals using bootstrap samples

Another challenge associated with sparse sampling of behavior and/or neuronal activity is estimating the reliability of tuning curves. One method to circumvent this problem is to derive statistics (mean and confidence interval) from bootstrapped samples (Kass et al., 2014). To this end, we can measure several times (e.g. *n* = 1000 samples) our likelihood P(A | S) on a portion of the data (e.g. 50% randomly chosen data points) and with replacement (the same data points can be used in different bootstrap samples; fig. 1q). Using these parameters (*n* = 1000 bootstrap samples using 50% of the data), we display every bootstrap tuning curve along with the corresponding 95% confidence interval (fig. 1r).

### 1.3 Information metrics

Numerous studies have applied metrics derived from information theory (Shannon, 1948) to neuronal data (Skaggs et al., 1993; see Dimitrov et al., 2011 for review). While the information held within a single spike is difficult to approximate with calcium imaging, mutual information (MI) can be used to describe the amount of information about one variable through observation of another variable (neuron activity and behavioral state in our example) using unconditional and joint probabilities:

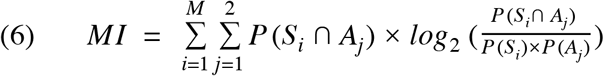

where *M* is the total number of possible behavioral states, *P* (*S_i_ ∩ A_j_*) is the joint probability of behavioral state *i* to be observed concurrently with activity level *j*. Note that in our case we consider only two levels of activity *j* (active versus inactive). The MI index is a practical way of sorting neurons by the amount of information they encode (supplementary fig. 1), and it was previously found that although related, MI is more reliable and scalable than other spatial information metrics (Souza et al., 2018). In our example, cell #4 displays 0.21 bits of information while considering only trajectories to the right (fig. 1j). On the other hand, it is possible to assess the significance of MI values by using the same techniques described in section 1.3. An example of MI values compared to shuffled surrogate can be found in supplementary fig. 1a.

### 2. Decoding of behavior from calcium imaging data

Extracting tuning curves for each individual neuron can shed light about their activity pattern but does not fully explain a particular behavior. Importantly, the temporal coordination of large neuronal assemblies is likely to provide more information about the specificity of generated behaviors. In our example, we would like to understand the relationship between location (without discriminating left/right trajectories at first) and the activity patterns of a large (~400) cell assembly. When a given neuron is active, the posterior probability density function (Eq. 5) computed earlier gives an estimate of the likely position of the mouse on the linear track. The most likely position of the mouse given that a neuron can be estimated as the location corresponding to the maximum value of the posterior probability density function. *P(S)* can be measured directly (in our case, it is the general likelihood of finding the mouse in any given location; fig. 2a, teal line in the bottom panel) or kept uniform. In the latter case, we make no prior assumption about the potential location of the mouse on the linear track and attribute equal probability for each location (fig. 2a, orange line in the bottom panel).

**Figure 2.**
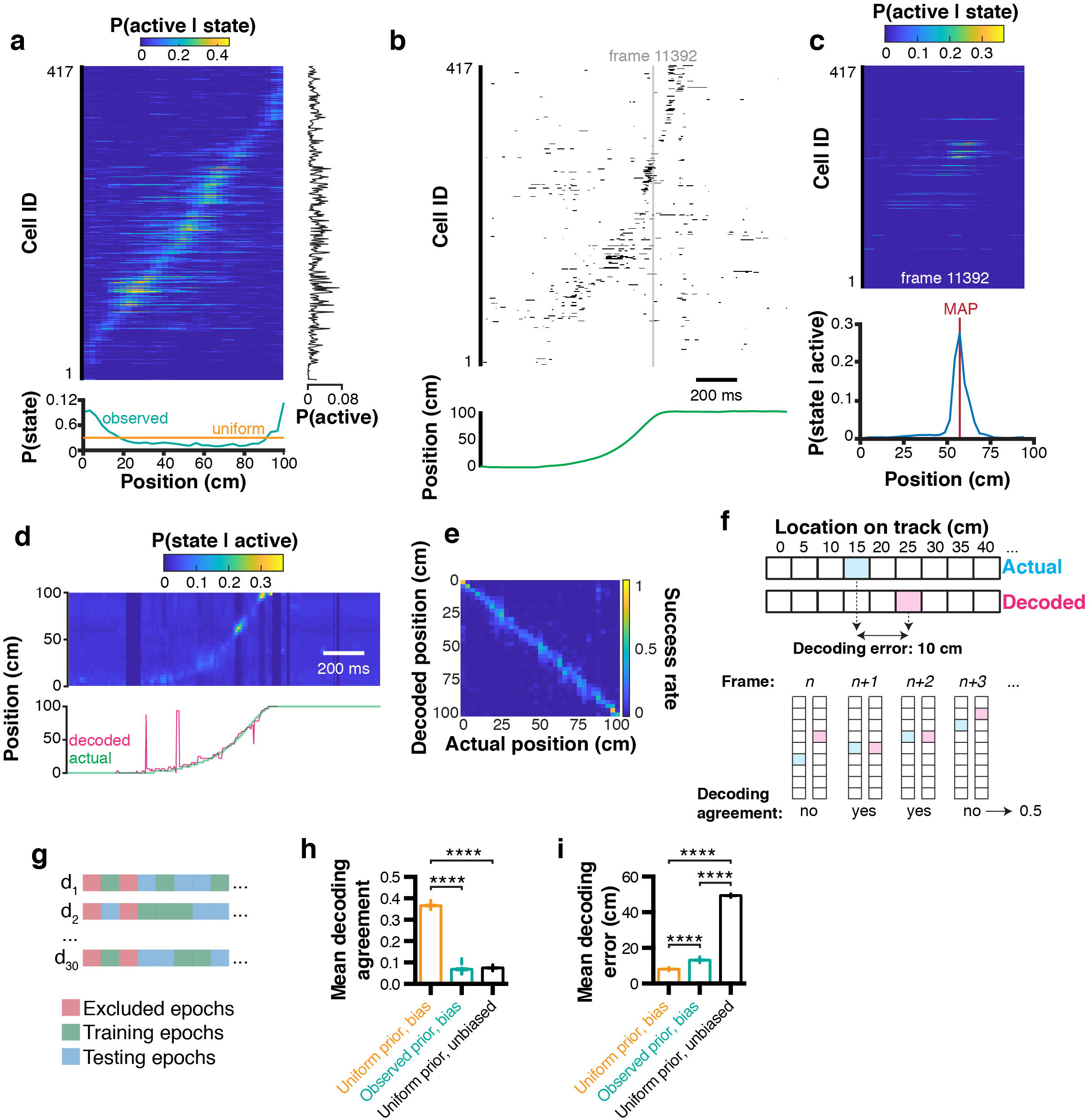
Bayesian decoding of behavior from calcium imaging recording. a, spatial tuning curves for each individual CA1 neuron (data sorted from location of peak probability P(active|state)), and corresponding marginal likelihood of being active (right-hand side), and prior probability of being in a given state (= location; bottom). b, raster plot of binarized cell activity and corresponding position on the linear track (bottom). c, tuning curves of cells corresponding to their state at frame 11392 (in b) and subsequent posterior probability of being in a location on the linear track given cell activity (bottom). Location was estimated using maximum *a posteriori* (MAP). d, posterior probabilities for each frame estimated from ongoing binarized calcium activity, and corresponding actual (green) and decoded (pink) location estimated with MAP. e, confusion matrix of actual vs decoded position. f, method used to compute Euclidean distance decoding error (top) and decoding agreement (bottom). g, paradigm used to train and test the decoder on different epochs of the dataset. h, effect of prior and bias (marginal likelihood of cell being active) on decoding agreement. i, same for decoding error. Color codes in a,c,d,e: dark blue = low probability, yellow = high probability.

#### 2.1. Decoding behavior using activity from multiple neurons

To predict the mouse state using the activity from multiple neurons, it is more efficient to take the product (rather than the sum) of *a posteriori* probabilities, because null values translate into an absolute certainty that the mouse cannot be in a given state considering the activity of a given neuron. Importantly, this can only be done under the assumption that every neuron is independent from each other, which is unlikely to be the case in practice because of neuroanatomical connectivity or other reasons: for example on a linear track the same sequence of neuronal activity is expected to be observed for each trajectory. In the case of interdependent neurons, posterior probabilities would have to be computed for each dependent neuronal ensemble, and the reconstructions would be expected to be more accurate at the expense of requiring significantly larger training datasets. Therefore, assuming that recorded neurons are independent trades off decoding accuracy for computing time. We can then rewrite our equation to include tuning curves from multiple neurons:

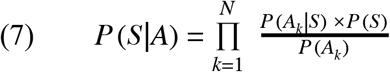

where *P* (*S*|*A*) is a vector of *a posteriori* behavioral states and *N* corresponds to the number of neurons used. In our example, we can measure the activity of every neuron *k* at a given time point (fig. 2b), derive the associated tuning curves (fig. 2c, top panel), and corresponding posterior location probability (fig. 2c, bottom panel). Importantly, while equation (7) is fundamentally correct, the repeated product of small values (such as probability values that are between 0 and 1) will lead to numerical underflow when computed on most softwares available currently. Although this is not a problem when decoding activity from a small number of cells, numerical underflow will prevent decoding activity from large sets of cell assemblies. One solution to this problem is to perform computations on a log scale. Additionally, using *exp(x)-1* and *log(1+x)* allows very small values to converge toward *x* instead of 0. Our equation can then take the form:

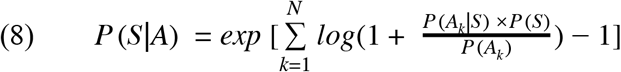

It is noteworthy that Eq. (7) and (8) are not formally equivalent. However, to reconstruct the position of the mouse, we will consider the location associated with the maximum *a posteriori* (MAP) which will remain the same value under such convex transform:

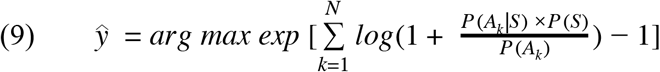

where *ŷ* is the estimated state among all states in S.

#### 2.2 Assessing decoding accuracy

In our example, we can compute the posterior probabilities for each individual timestep based on neuronal activity, and compare the actual versus decoded location on the linear track (fig. 2d). To visualize which states are associated with better/worse decoding error, we can compute a confusion matrix, which expresses the portion of time points where the actual state was successfully decoded (fig. 2e). This representation is also useful to identify which states are more difficult to decode. While confusion matrices are useful, they are not practical when it comes to summarizing decoding accuracy for large datasets and performing statistical analyzes. We thus propose two metrics: (1) decoding agreement, and (2) decoding error. We define decoding agreement as the portion of time where the exact state of the mouse was successfully decoded:

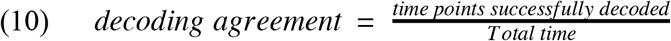

Therefore, decoding agreement is a value between 0 and 1. For instance, a value of 0.5 means that 50 % of time points have been successfully decoded. This approach is quite conservative: when the decoded state is only one bin away from the actual behavioral state, it would lead to a null decoding agreement while still being close to reality. Therefore, another metric commonly used in decoding analyzes is decoding error, which is the distance between the decoded behavior and the actual behavior. Note that in our case, the distance is explicitly Euclidean and can be expressed in cm. For one-dimensional data, equation (16) can be used to compute decoding error:

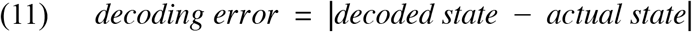

The decoding error may or may not be useful depending on the variables under study. For instance, in the case of auditory stimuli, the distance between tone frequencies might not necessarily be as meaningful as an actual spatial distance, as it is the case in our example. Moreover, its computation will be different for two-dimensional space, or head orientation, to list a few of the variables commonly studied. Importantly, to assess decoding accuracy, it is recommended not to test the decoder on epochs that were used to train the Bayesian decoder. Some epochs, such as periods of immobility in our case, can be excluded for both training and testing altogether. We propose here to train and test our decoder on non-overlapping sets of random epochs, repeat the process several times, and compute the average decoding agreement and decoding error (fig. 2g). Using this approach, we found in our conditions that using a uniform prior *P(S)* led to higher decoding agreement (0.37 ± 0.002, n = 30 trials; data expressed as mean ± SEM) compared to using observed prior probabilities (0.07 ± 0.004, n = 30 independent trials), or replacing the marginal likelihood (bias) *P(A)* by 1 (condition which we term ‘unbiased’ here; 0.07 ± 0.001, n = 30 independent trials; 1ANOVA, F_(2, 87)_ = 4521, P < 0.0001; fig. 2h). Similarly, decoding error was lower using a uniform prior (8.12 ± 0.08 cm, n = 30 independent trials) compared to using an observed prior (13.18 ± 0.15 cm, n = 30 independent trials) or in unbiased conditions (49.34 ± 0.08 cm, n = 30 trials; F_(2,87)_ = 44710, P < 0.0001; fig. 2i).

#### 2.3. Adding temporal constraints

Although decoding can be performed for each individual time point, this temporal independence can lead to spurious results (see decoded trajectory in fig. 2d, pink line in the bottom panel). Rationaly, if the mouse is at one end of the linear track, it is extremely unlikely to be found at the opposite end on the next frame. There are several ways to solve this problem and improve state estimation. A first method could be to build a transition matrix (such as one that would be used in a Markov process), and attribute null values to impossible transitions (such as going from one end of the linear track to the other), as well as uniform probabilities to adjacent states. One could then replace the observed or uniform prior *P(S)* by the appropriate realistic transition values at each individual timestep. Another method consists of taking the product of temporally adjacent posteriors. In that case, we would decode the mouse state by rewriting equation (7):

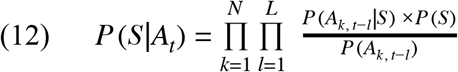

where *P* (*S*|*A_t_*) is the *a posteriori* distribution of states at time *t*. The number of past timesteps *l* used to estimate the mouse state at time *t* is determined by *L*. The effect of different values of *L* will be detailed in the following section. The advantage of this method is that it does not require to determine transition probabilities experimentally. In our conditions, temporal filtering can greatly improve reconstruction and remove erratic ‘jumps’ that can sometimes occur during decoding (fig. 3a).

**Figure 3.**
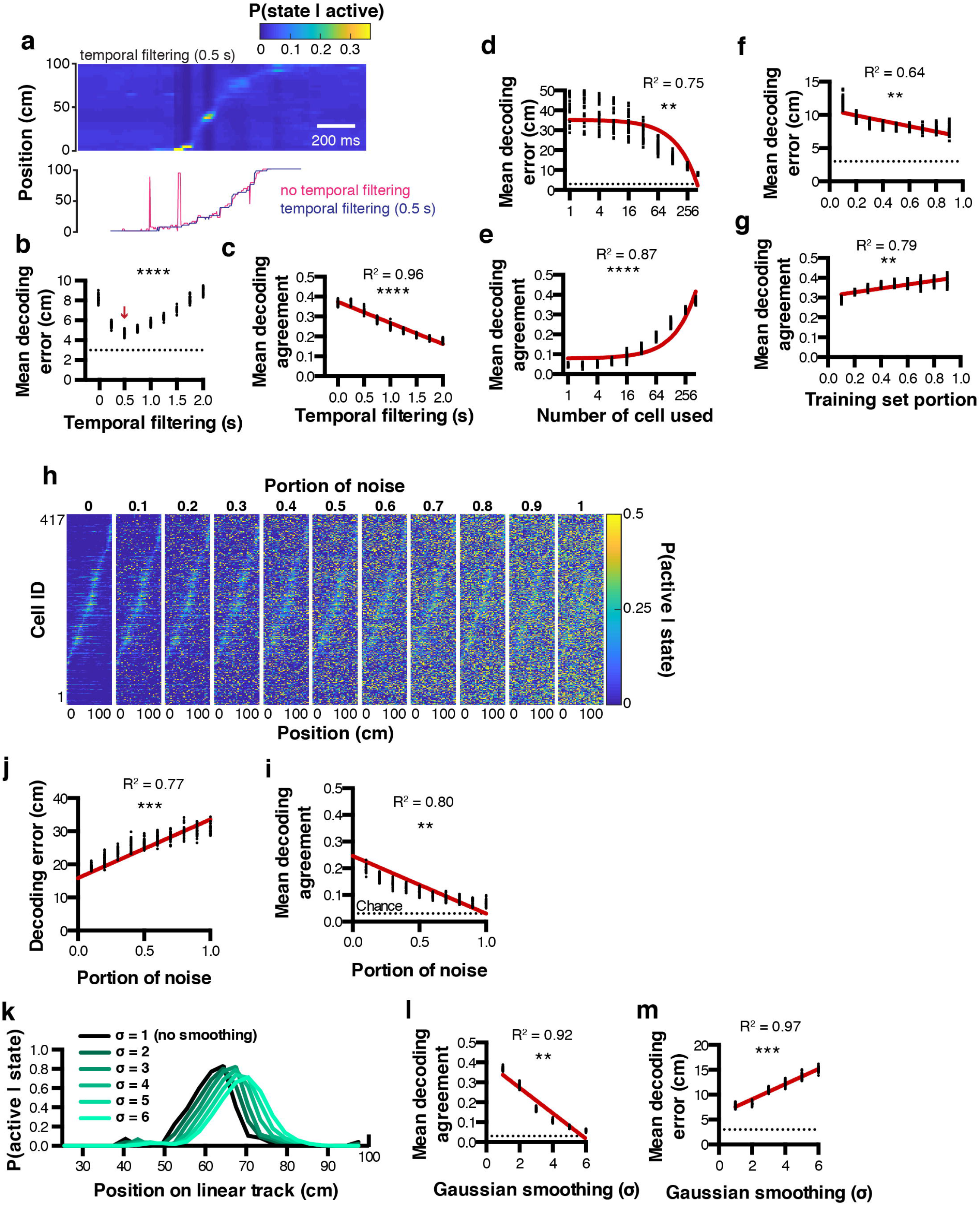
Decoding parameter estimation. a, example posterior probabilities P(state|active) when using a 0.5 s temporal filtering window (top), and corresponding decoded location estimated from MAP (bottom). b & c, effect of temporal filtering window size on decoding error and agreement, respectively. d & e, effect of the number of cells used on decoding error and agreement, respectively. f & g, effect of training set portion on decoding error and agreement, respectively. h, effect of random noise on spatial tuning curves. i & j, corresponding decoding agreement and error, respectively. k, effect of gaussian smoothing on spatial tuning curves. l & m, corresponding decoding agreement and error, respectively. Color codes in a & h,: dark blue = low probability, yellow = high probability.

#### 2.4. Parameter optimization

To find the best conditions to decode neuronal activity, it is possible to optimize parameters including (but not restricted to): number of cells used, portion of training set, temporal filtering window, or spatial filtering of tuning curves. For instance, we performed decoding on 30 sets of random epochs using several temporal filtering window sizes ranging from 0 (no filtering) to 2 s, and found that better reconstructions could be achieved using a 0.5 s filtering window, leading to smaller decoding errors (4.73 ± 0.04 cm, n = 30 independent trials per filtering window; 1ANOVA, F_(8,261)_ = 1002, P < 0.0001; fig. 3b). Interestingly, the bigger the temporal filtering window, the lower the decoding agreement (Pearson correlation, R^2^ = 0.96, P < 0.0001, n = 30 independent trials per filtering window; fig. 3c). As expected, considering more cells during the reconstruction resulted in decreased decoding error (Pearson correlation, R^2^ = 0.75, P < 0.0012, n = 30 independent trials per cell number; fig. 3d) and increased decoding agreement (Pearson correlation, R^2^ = 0.87, P < 0.0001, n = 30 independent trials per cell number; fig. 3e). We also tested the influence of the training/testing length ratio on reconstruction accuracy and found that good reconstruction can be achieved by using testing epochs that represent beyond 30 % of the total experiment length. Mean decoding error decreased as the training set portion increased (Pearson correlation, R^2^ = 0.64, P = 0.01, n = 30 independent trials per training set portion tested; fig. 3f), while mean decoding agreement increased (Pearson correlation, R^2^ = 0.79, P = 0.0013, n = 30 independent trials per training set portion tested; fig. 3g). We next assessed the robustness of tuning curves to random noise. To this end, we computed tuning curves as described previously, then replaced a portion (between 0 and 1, with 0.1 incremental steps) of the tuning curves data with random probability values (fig. 3h). Addition of noise was correlated with decreased decoding agreement (Pearson correlation, R^2^ = 0.80, P = 0.0014, n = 30 independent trials per noise amount; fig. 3i), and increased decoding error (Pearson correlation, R^2^ = 0.77, P = 0.0004, n = 30 independent trials per noise amount; fig. 3j). Finally, we tested the impact of smoothing tuning curves on decoding accuracy. Gaussian smoothing is often performed in the context of place cell studies, presumably to improve the visualization of assumed place fields (O’Keefe and Burgess, 1996; Hetherington and Shapiro, 1997). In our conditions, we found that Gaussian smoothing of tuning curves was systematically associated with decreased decoding agreement (Pearson correlation, R^2^ = 0.92, P = 0.0025; n = 30 independent trials per gaussian sigma value; fig. 3l), together with increasing decoding error (Pearson correlation, R^2^ = 0.97, P = 0.0003, n = 30 independent trials per gaussian sigma value; fig. 3m).

#### 2.5. Optimal method to binarize neuronal activity

In our conditions, we used a simple binarizing algorithm that transformed rising periods of calcium transients into periods of activity. We compared this method to a simple z-score threshold where all activity above 2 standard-deviations is considered active, and to a deconvolved signal using a first order autoregressive model (see methods), where all values above zero are considered as periods of activity. To quantify the accuracy of these methods, we performed *in vitro* electrophysiological recordings in the cell attached configuration, in conjunction with 1-photon calcium imaging (supplementary fig. 2a). We extracted calcium transients from the recorded cell (supplementary fig. 2b) so as to contrast these signals with ground truth spiking activity (supplementary fig. 2c). Interestingly, calcium transients appeared much longer in these conditions, and our binarizing method only matched the later portion of transients rising periods (supplementary fig. 2d). In the following analyses, we seperated the portion of action potentials successfully labeled as active periods in the corresponding calcium imaging recording frame, as well as the portion of total recorded frames that were either correctly labeled as active if they were associated with at least one action potential, or inactive if they did not. Using a deconvolved trace to estimate neuronal activity resulted in a higher number of action potentials successfully detected as corresponding calcium imaging active periods (0.94 ± 0.032) compared to our binarizing algorithm (0.49 ± 0.067) or a simple z-score threshold (0.65 ± 0.075; 1ANOVA, F_(2, 108)_ = 13.71, P < 0.0001, n = 37 detection windows; supplementary fig. 2e). Furthermore, both the portion of true negatives (calcium imaging frames binarized as inactive that indeed contained no action potential divided by the total number of recorded frames) and the portion of true positives (calcium imaging frames binarized as active that indeed contained at least one action potential, divided by total number of recorded frames) were comparable between methods (supplementary fig. 2f & g respectively).

Interestingly, these *in vitro* results did not compare to our *in vivo* conditions. When computing tuning curves for the neuron presented in fig. 1, using a simple threshold resulted in a larger place field, while binarizing data from a deconvolved trace resulted in two peaks (supplementary fig. 3a). While there is no ground truth data to conclude which method is best to compute tuning curves, decoding analyzes can shed a light on this question, because animal behavior can be used as ground truth data (the higher the decoding accuracy, the closer to ground truth). We thus trained our decoder (Eq. 9) using tuning curves computed from binarized activity derived using a simple z-score threshold, a deconvolved trace, or using our binarizing method. We found that using both our binarizing method (4.74 ± 0.0039 cm) or a deconvolved trace (4.81 ± 0.048 cm) led to lower decoding errors compared to using a simple threshold (5.18 ± 0.051 cm, F_(2,87)_ = 26.22, P < 0.0001, n = 30 independent trials for each binarizing method.)

#### 2.6. Decoding two-dimensional behavioral variables

The decoding method presented above is scalable to a large variety of behaviors. However, sometimes it can be useful to represent behaviors in more than one dimension. This is for instance the case with spatial location in larger environments. We will now show that the principles presented above can easily be translated to more dimensions. To this end, we recorded neuronal activity using calcium imaging in a mouse exploring an open-field for the first time. Calcium transients are then extracted and binarized, along with the x and y mouse position (fig. 4a). It is possible to plot periods of activity of one cell in 2D space, and color code this activity to visualize the stability of such activity in time/space (fig. 4b). Relative occupancy (fig. 4c) and P(active|state) probabilities can be computed for each state (3 x 3 cm spatial bin) the same way as presented above (fig. 4d). To assess the confidence of such result, it is also possible to shuffle data circularly and compute P(active|state) probability maps (fig. 4e). From these shuffled probability maps, we can derive the level of significance using a p-value that corresponds to the number of shuffled data points above the actual data divided by the number of shuffled surrogates (fig. 4g). We can then derive a thresholded place field that only retains significant values of P(active|state) (fig. 4h). Importantly, it is noteworthy that the standard-deviation of the shuffled distribution (fig. 4i) is negatively correlated to the relative occupancy (Pearson correlation, R^2^ = 0.47, P < 0.0001; fig. 4j). This suggests that for states with high P(active|state) probabilities, significance can be higher if the state displays high occupancy, and lower if the state displays low occupancy. We also assessed the effect of temporal filtering on the quality of the reconstructions and found that in our conditions, a 1.6 s filtering window yielded best results (1ANOVA, F_(39,1160)_ = 72.31, P < 0.0001, n = 30 independent trials per temporal filter window size; fig. 4k). As for one-dimensional data, Gaussian filtering of tuning maps (2D tuning curves) consistently increased the decoding error (Pearson correlation, R^2^ = 0.99, P < 0.0001, n = 30 independent trials per Gaussian sigma value; fig. 4l).

**Figure 4.**
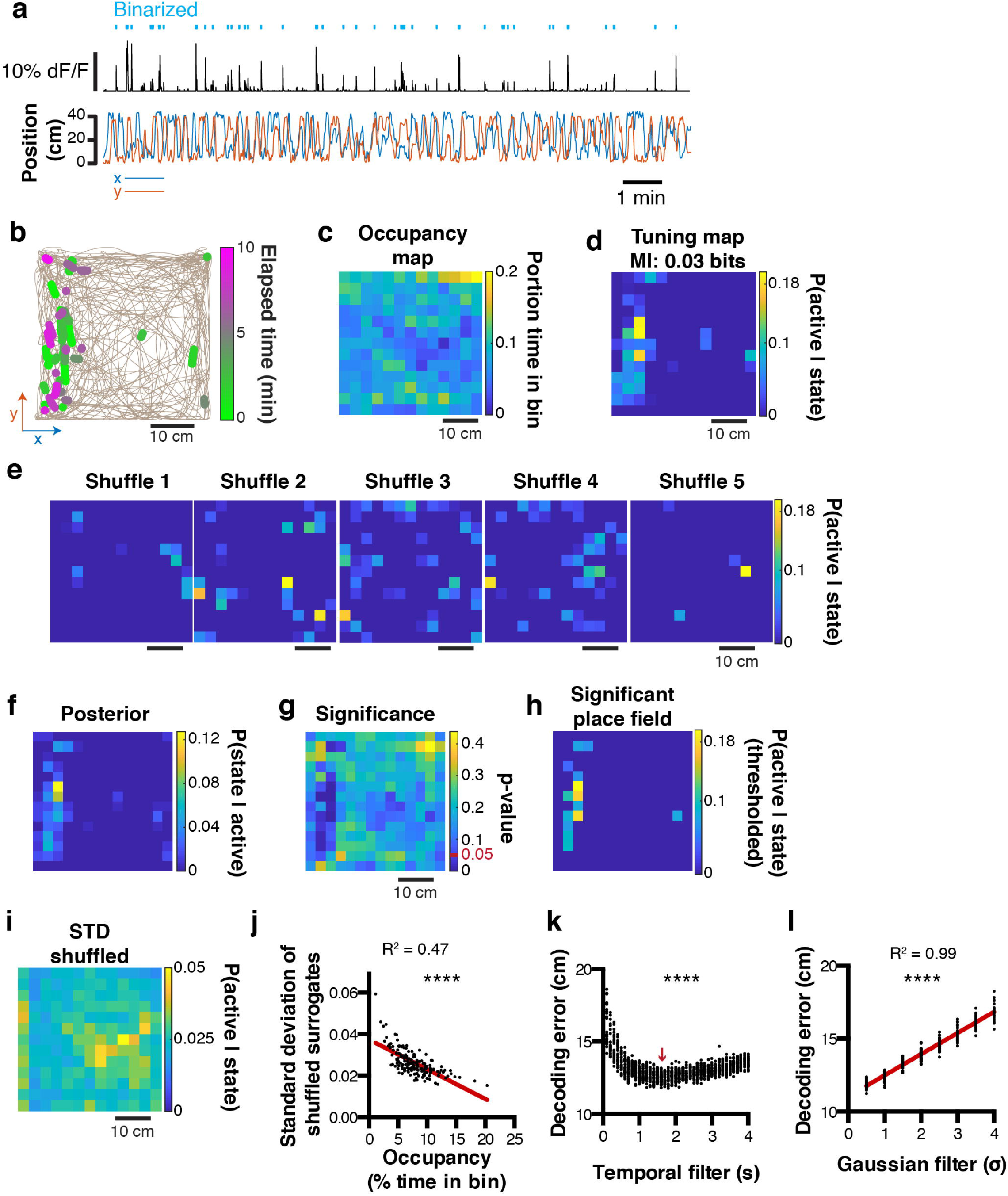
Decoding two-dimensional behaviors. a, *x,y* coordinates of mouse location in an open field (bottom) and corresponding raw calcium trace of one example cell and binarized activity (top). b, top view of mouse trajectory (beige trace) with overlaid location corresponding to neuronal activity (green, early activity; magenta, late activity). c, relative occupancy in the open field. d, tuning map (probability of being active given location) of one neuron. Top, corresponding MI index. e, example tuning maps computed from shuffled calcium traces. e, posterior probability P(state|active) for the same cell. g, p-value computed from the actual tuning map and corresponding shuffled surrogates. h, thresholded place field using only significant P(active|state) values. i, standard-deviation of the shuffled distribution. j, scatter plot comparing the standard-deviation of the shuffled distribution, and the mouse open field occupancy. k, effect of temporal filtering on decoding error in the open field. The red arrow indicates the temporal filtering window size yielding the lower decoding error. l, effect of gaussian smoothing of tuning maps on decoding error in the open field.

### 3. Refining encoding predictions

The ultimate goal of decoding neuronal activity is to improve our understanding of the relationship that may exist between neurons and behavior. In other terms, in addition to predicting behavior from neuronal activity, one should be able to predict neuronal activity from behavior. These encoding predictions can easily be derived from equation (3) since *P* (*A*|*S*) represents the likelihood of a given neuron to be active while an animal is in a given state. This probability distribution can be interpreted as both a direct measure from our data, and a prediction of activity rate as a function of the animal location. Consequently, if *P* (*A*|*S_i_*) would take the value 1 for the given states *i*, we would be absolutely certain that the considered neuron would be active in these given states. In our linear track example, we can refine our encoding prediction since it has been previously shown that hippocampal place cell activity on a linear track tend to be unidirectional: some cell will fire in one given location, but only when being traversed in one direction (McNaughton et al., 1983; Markus et al., 1995). If the peak probability of being active *P* (*A*|*S*) for a neuron that displays a prominent place field is only 0.5, it could be due to the fact that this cell fires only 50 % of the time, when the animal is running in one direction, but not in the other. We will now demonstrate that it is possible to predict neuronal activity from estimated tuning curves, and that refining our behavioral states by including directional information can increase our encoding prediction, i.e. the confidence we have that a neuron will fire given that the animal is in a given state. To this end, we extracted tuning curves from neurons being active on the linear track using either location only (fig. 5a), or by considering both location and direction (fig. 5b). Note that using the later method, the peak probability *P* (*A*|*S*) greatly increases. When comparing probability values obtained from the same cells but using either technique, it seems that most cells’ *P* (*A*|*S*) increase when including traveling direction (Pearson correlation, r = 0.8781, R^2^ = 0.77, P < 0.0001; fig. 5c). Interestingly, this is not the case for a minority of cells, indicating that some cells still display preferential activity for absolute location, regardless of direction. We then reconstructed neuronal activity using the *P* (*A*|*S_i_*) of each neuron for each known state *i* (location in our example) of the mouse. For display purposes, we only show probability values greater than 0.5 (fig. 5d). To estimate the confidence in neuronal activity predictions, we can use the same bootstrap method presented earlier to build 95% confidence intervals for *P*(*A*|*S*) tuning curves computed by considering left, right, or both traveling directions (fig. 5e; n = 1000 bootstrap samples). The advantage of this method is that it gives a clear range of prediction accuracy that is easily interpretable. In our example neuron #4, it is apparent that greater encoding predictions can be achieved when only considering travels to the right (fig. 5e). Furthermore, we can use mutual information as a measure of how much uncertainty about neuronal activity can be reduced by knowing the state of the animal. In our example, we found that MI values were the highest when only considering travels to the right (0.22 ± 0.0005 bits), followed by considering both directions (0.09 ± 0.0002 bits), and only travels to the left (0.02 ± 0.0001 bits; 1ANOVA, F_(2, 2997)_ = 44.84, P < 0.0001; n = 1000 bootstrap samples; fig. 5f).

**Figure 5.**
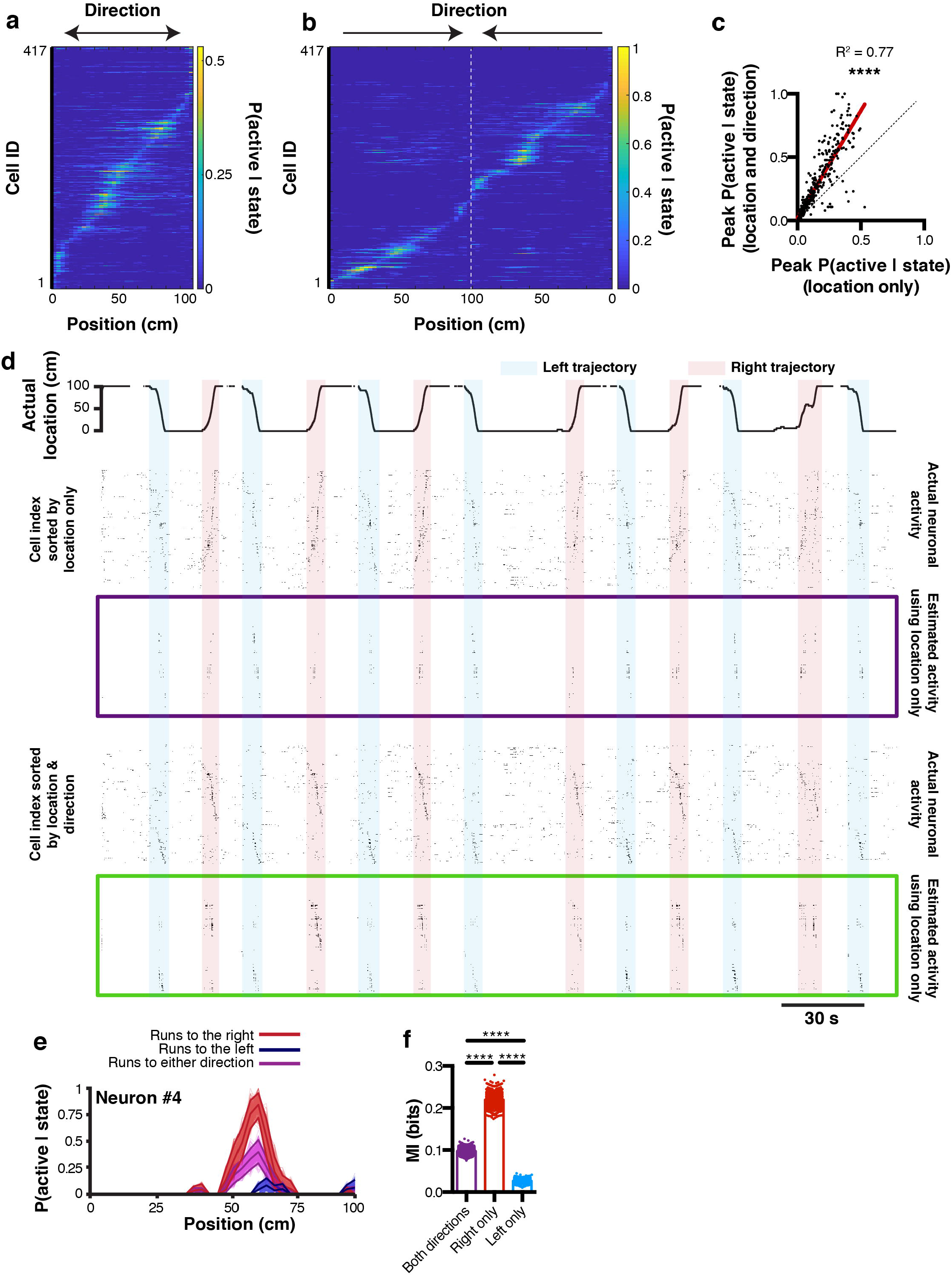
Reconstructing neuronal activity and refining tuning curves. a, tuning curves of every neuron sorted by peak activity in the linear track. b, same, but after discriminating left and right trajectories. c, relationship between peak P(active|state) likelihood computed using either method (location only versus location and direction). d, actual location of the mouse in the linear track (top), and corresponding actual and reconstructed neuronal activity using location only (purple box), as well as actual and reconstructed neuronal activity using both location and direction (green box). e, P(active|state) computed using either right (red), left (blue), or both (purple) trajectories. Thickness of curves indicate 95% confidence interval computed from n = 1000 bootstrap samples. f, corresponding MI for each bootstrap sample.

## Discussion

We show here that representing neuronal activity extracted from calcium imaging data by a binary state (active vs inactive) is sufficient to approximate the state of a neuronal assembly. While such binarization was previously proposed as an intermediate step to perform decoding (Ziv et al., 2013), here we generalize this principle and propose several additional metrics to describe the strength of neuronal tuning to behavioral variables. In particular, several methods can be used to binarize calcium activity, but because the rise time of calcium transients contains the vast majority of action potentials, binarizing methods should aim at labeling specifically these epochs as periods of activity. Importantly, optimizing methods and parameters using *in vitro* conditions cannot necessarily be translated to data acquired *in vivo* because calcium transients differ fundamentally across conditions, even if most variables are the same (animal strain/age, viral construct and dilution).

Information on neuronal coding can be extracted using simple methods and minimal data processing. Importantly, three metrics can be used to describe neurons: the likelihood of being active in a given behavioral state P(active|state), the posterior probability of the animal being in a state given neuronal activity P(state|active), and mutual information - a measure of reduction in uncertainty of the mouse state that results from knowing neuronal activity, and vice versa. Furthermore, we propose to determine significance by either computing a p-value value derived from shuffled surrogates, or estimating confidence intervals using bootstrap resampling. In particular, the later method provides confidence in encoding predictions that are easily interpretable.

While action potentials recorded with electrophysiological techniques constitute discrete temporal events, calcium imaging signals take the form of a continuous vector which prevents the direct computation of previously used information metrics: bits/s and bits/spike. On the other hand, MI values can easily be derived from probabilities computed above and provide useful insights in the amount of uncertainty that is reduced about the animal state given neuronal activity. It is noteworthy that the MI is sensitive to the number of bins used, therefore faithful comparisons between electrophysiological and calcium imaging data would require to compute MI values on electrophysiological and calcium imaging data where neuronal activity was binarized the same way as well as behavior discretized using the same binning parameters. Furthermore, while bursting activity is associated with large calcium transients that result in good decoding accuracy, single spikes might lead to changes in GCamp6 fluorescence that are too small to be detected, and could be associated with larger decoding errors in some conditions.

Unsurprisingly, we found that MI values are significantly correlated with peak P(active|state) probabilities, while in parallel, behavioral states with low occupancy display more variability in the surrogate data, which indicates that higher significance can be achieved when the sampling of behavioral states is higher. Importantly, the choice of a null hypothesis should be considered carefully. On one hand, if it is hypothesized that neurons become active in response to an external stimulus, then one should permute neuronal activity in respect to the time course of external stimuli under the null hypothesis. On the other hand, if it is hypothesized that the temporal organization of neurons (e.g. sequences) underlies certain cognitive processes (e.g. replay), then permutations should be performed so as to remove temporal patterns of neuronal activity (i.e. shuffle each neuron activity vector independently) under the null hypothesis.

Using such a probabilistic approach allows to derive predictions based on Bayes theorem. In our conditions, we found that minimal *a priori* (uniform distribution of states likelihood) yielded better results. Adding temporal constraints could decrease decoding error but not decoding agreement. Consequently, these filters have to be optimized on a case by case basis depending on the goal of the study. Interestingly, smoothing probability distributions had negative effects in our conditions, most likely due to the asymmetric nature of place fields when unfiltered. Such post-processing methods thus have to be used with caution, and while they can improve the visualization of particular phenomena such as place fields, they can result in spurious interpretations. While the method we describe here might not apply to any type of behavior, we present examples of one- and two-dimensional datasets, and the number of dimensions being studied should not be a limiting factor. One of the most significant limitations with this method arises directly from behavioral sampling, and only behaviors that are directly sampled can be reconstructed or used to predict neuronal activity. For alternative approaches that do not make assumptions about behaviors being encoded in neuronal activity, the use of dimensionality reduction techniques has been proposed (Rubin et al., 2019), and if sequential patterns are hypothesized to underlie behaviors, matrix factorization methods can be used instead (Mackevicius et al., 2018). Moreover, when multiple variables contribute to variability in neuronal activity, generalized linear models can outperform the methods presented here (Park et al., 2014), at the expense of requiring significantly more data points which may or may not be compatible with calcium imaging approaches, depending on the activity rate of neurons being studied.

Finally, we propose a simple method to characterize neuronal encoding and predict neuronal activity. This method is useful in refining the behavioral components that can determine neuronal activity. As such, the quality of models that can be drawn from observations largely depends on the very nature and accuracy of these observations. In particular, increasing the amount of information concerning a certain behavior can result in a refinement of the underlying model of neuronal activity. Perfect predictions of neuronal activity on the simple basis of behavior is a difficult endeavor however, because such activity is not only determined by external variables (behavior) but also internal variables including animal state, and pre-synaptic activity that is often inaccessible to the observer. In this context, previous work has outlined organized patterns of neuronal activity that are usually associated with spatial location while animals did not perceive any external stimuli other than self-motion (Villette et al., 2015). Moreover, taking into account the independence of neuronal activity could also improve the quality of predictions. In particular, including pairwise (Pillow et al., 2008; Meshulam et al., 2017) or temporal (Naud and Gerstner, 2012) correlations of neuronal activity could reduce decoding error. Note that these temporal correlations would also be taken into account when using long short-term memory (LSTM) artificial neural networks (Tampuu et al., 2019), thus increasing reconstruction accuracy at the expense of interpretability. Rather than proposing a sophisticated analysis pipeline, the methods presented here have the advantage of remaining simple, requiring only few data points, and are easily interpretable using metrics that can facilitate the communication of results along with significance and confidence intervals, making it an appropriate tool for exploration of calcium imaging data in conjunction with behavior.

## Materials and methods

### Surgical procedures

All procedures were approved by the McGill University Animal Care Committee and the Canadian Council on Animal Care. For the linear track and open field data, one adult mouse (~2 months) was anesthetized with isoflurane (5% induction, 0.5-2% maintenance) and placed in a stereotaxic frame (Stoelting). The skull was completely cleared of all connective tissue, and a ~500 μm hole was drilled. We then injected the AAV5.CamKII.GCaMP6f.WPRE.SV40 virus (Addgene # 100834; 200 nL at 1 nl.s^-1^) in hippocampal CA1 using the following coordinates: anteroposterior (AP) −1.86 mm from bregma, mediolateral (ML) 1.5 mm, dorsoventral (DV) 1.5 mm. 2 weeks following the injection, the mouse was anesthetized with isoflurane and the skull was cleared. A ~2 mm diameter hole was perforated in the skull above the injection site. An anchor screw was placed on the posterior plate above the cerebellum. The dura was removed, and the portion of the cortex above the injection site was aspirated using a vacuum pump, until the corpus callosum was visible. These fiber bundles were then gently aspirated without applying pressure on the underlying hippocampus, and a 1.8 mm diameter gradient index (GRIN; Edmund Optics) lens was lower at the following coordinates: AP −1.86 mm from bregma, ML 1.5 mm, DV 1.2 mm. The GRIN lens was permanently attached to the skull using C&B-Metabond (Patterson dental), and Kwik-Sil (World Precision Instruments) silicone adhesive was placed on the GRIN to protect it. 4 weeks later, the silicone cap was removed and CA1 was imaged using a miniscope mounted with an aluminium base plate while the mouse was under light anesthesia (~0.5 % isoflurane) to allow the visualization of cell activity. When a satisfying field of view was found (large neuronal assembly, visible landmarks), the base plate was cemented above the GRIN lens, the miniscope was removed, and a plastic cap was placed on the base plate to protect the GRIN lens.

### Behavior and miniscope recordings

After baseplating, the mouse was gently handled for ~5 min per day for 7 days. The mouse was then water-scheduled (2 h access per day), and placed on a 1 m long linear track for 15 min. 10% sucrose in water rewards were placed at each end of the linear track, and the mouse had to consume one reward before getting the next one delivered. Miniscope recordings were performed at 30 Hz for 15 min every day, and decoding was performed on the last training day (day 7). The following week, the mouse was allowed to freely explore for 15 min a 45 x 45 cm dark gray open-field that contained visual cues, and miniscope recordings were performed at 30 Hz for the entire duration of the exploration (15 min).

### Miniscope and behavior video acquisition

Miniscopes were manufactured using open source plans available on www.miniscope.org and as described previously (Ghosh et al., 2011; Cai et al., 2016; Aharoni and Hoogland, 2019). Imaging data was acquired using a CMOS imaging sensor (Aptina, MT9V032) and multiplexed through a lightweight coaxial cable. Data was acquired using a data acquisition (DAQ) box connected via a USB host controller (Cypress, CYUSB3013). Data was recorded using a custom written acquisition software relying on Open Computer Vision (OpenCV) librairies. Video streams were recorded at ~30 frames per second (30 Hz) and saved as uncompressed.avi files. Animal behavior was recorded using a webcam, and the DAQ software simultaneously recorded timestamps for both the miniscope and behavior camera in order to perform subsequent alignment.

### Calcium imaging analysis

Calcium imaging videos were analyzed using the MiniscopeAnalysis pipeline (https://github.com/etterguillaume/MiniscopeAnalysis). In particular, we first applied rigid motion correction using NoRMCorre (Pnevmatikakis and Giovannucci, 2017). 1000 frame videos were then concatenated into a large video file after a 2 fold spatial downsampling. Spatial components as well as calcium traces were then extracted using CNMFe (Zhou et al., 2018) using the following parameters: gSig = 3 pixels (width of gaussian kernel), gSiz = 15 pixels (approximate neuron diameter), background_model = ‘ring’, spatial_algorithm = ‘hals’, min_corr = 0.8 (minimum pixel correlation threshold), min_PNR = 8 (minimum peak-to-noise ratio threshold). When applicable, calcium traces were deconvolved with OASIS (Friedrich et al., 2017), using an autoregressive model with order p = 1 and using the ‘constrained’ method.

### *In vitro* patch-clamp electrophysiology

One adult mouse (~ 2 months) was stereotaxically injected with a GCaMP6f construct (AAV5.CamKII.GCaMP6f.WPRE.SV40 virus, Addgene # 100834; 0.4 μL at 0.06 μl/min) in hippocampal CA1. 2 weeks later, it was deeply anesthetized using ketamine/xylazine/acepromazine mix (100, 16, 3 mg/kg, respectively, intraperitoneal injection), and intracardially perfused with cold N-methyl-d-glutamine (NMDG) recovery solution (4° C), oxygenated with carbogen (5% CO2 / 95% O2). The NMDG solution contained the following (in mM): 93 NMDG, 93 HCl, 2.5 KCl, 1.2 NaH2PO4, 30 NaHCO3, 20 HEPES, 25 glucose, 5 sodium ascorbate, 2 thiourea, 3 sodium pyruvate, pH adjusted to 7.4 with HCl before adding 10 MgSO_4_ and 0.5 CaCl_2_. Following NMDG perfusion, brains were quickly removed and immersed for an additional 1 minute in cold NMDG recovery solution. Coronal slices (300 μm) were cut using a vibratome (Leica-VT1000S), then collected in a 32°C NMDG recovery solution for 12 minutes. Slices were transferred to room temperature and oxygenated artificial cerebrospinal fluid (aCSF) containing the following (in mM): 124 NaCl, 24 NaHCO3, 2.5 KCl, 1.2 NaH2PO4, 2 MgSO4, 5 HEPES, 2 CaCl_2_ and 12.5 glucose (pH 7.4). Patch pipettes (3-5 MΩ) were filled with internal solution, containing the following (in mM): 140 K gluconate, 2 MgCl2, 10 HEPES, 0.2 EGTA, 2 NaCl, 2 mM Na2-ATP and 0.3 mM Na2-GTP, pH adjusted to 7.3 with KOH, 290 mOsm. Slices were transferred to a submerged recording chamber filled with aCSF (2-3 ml /min flow rate, 30 °C), continuously oxygenated with carbogen. All reagents were purchased from Sigma-Aldrich, unless stated otherwise. Extracellular cell-attached patch-clamp recordings were used for monitoring spontaneous cell firing activity from hippocampal pyramidal neurons expressing GcAMP6f (identified under EGFP-fluorescence). The recording pipette was held at a potential of −70 mV Imaging of GcAMP6f-expressing pyramidal cells was performed with a 60x Olympus water immersion objective (LUMPLFLN60X/W, NA 1.0) and acquired at 10 Hz using Olympus cellSens software. Electrophysiological signals were amplified, using a Multiclamp 700B patch-clamp amplifier (Axon Instruments), sampled at 20 kHz, and filtered at 10 kHz.

### Statistics

Statistical analyses were performed using GraphPad Prism version 6.00 (GraphPad Software, La Jolla, California USA). All data are presented as mean ± standard error of the mean (SEM) and statistical test details are described in the corresponding results. All t-tests are two-tailed. Normality distribution of each group was assessed using the Shapiro-Wilk normality test and parametric tests were used only when distributions were found normal (non-parametric tests are described where applicable). 1ANOVA: one-way ANOVA; 2ANOVA: two-way ANOVA; RMANOVA: repeated measure ANOVA. *p* < 0.05 was considered statistically significant. *, *p* < 0.05; **, *p* < 0.01; ***, *p* < 0.001, ****, *p* < 0.0001.

## Code and data availability

All the code and data presented here can be downloaded at the following address: https://github.com/etterguillaume/CaImDecoding

## Acknowledgments

We thank Ke Cui for support maintaining the colony and perfusing mice, Daniel Aharoni for support using the miniaturized fluorescence microscopes (www.miniscope.org), Bruno Rivard for manufacturing the miniscopes, and Suzanne van der Veldt for helpful comments on the manuscript. This work is supported by Brain Canada, Fonds de la recherche en Santé du Québec (FRSQ), the Canadian Institutes of Health Research (CIHR), the Natural Sciences and Engineering Research Council of Canada (NSERC), and the Alzheimer Society of Canada.

## Contributions

GE and SW designed the study. GE performed surgeries, behavioral experiments, and analyzes. FM performed patch clamp *in vitro* electrophysiology. GE and SW wrote the manuscript.

## Competing interests

The authors declare no competing interests.

## Notes

#### Summary of Updates

- Errors in terminology have been corrected (prior vs likelihood in Bayesian framework) - Corrected minor mistakes and typos

