## Supplementary information for "A probabilistic framework for decoding behavior from in vivo calcium imaging data"

Etter et al.

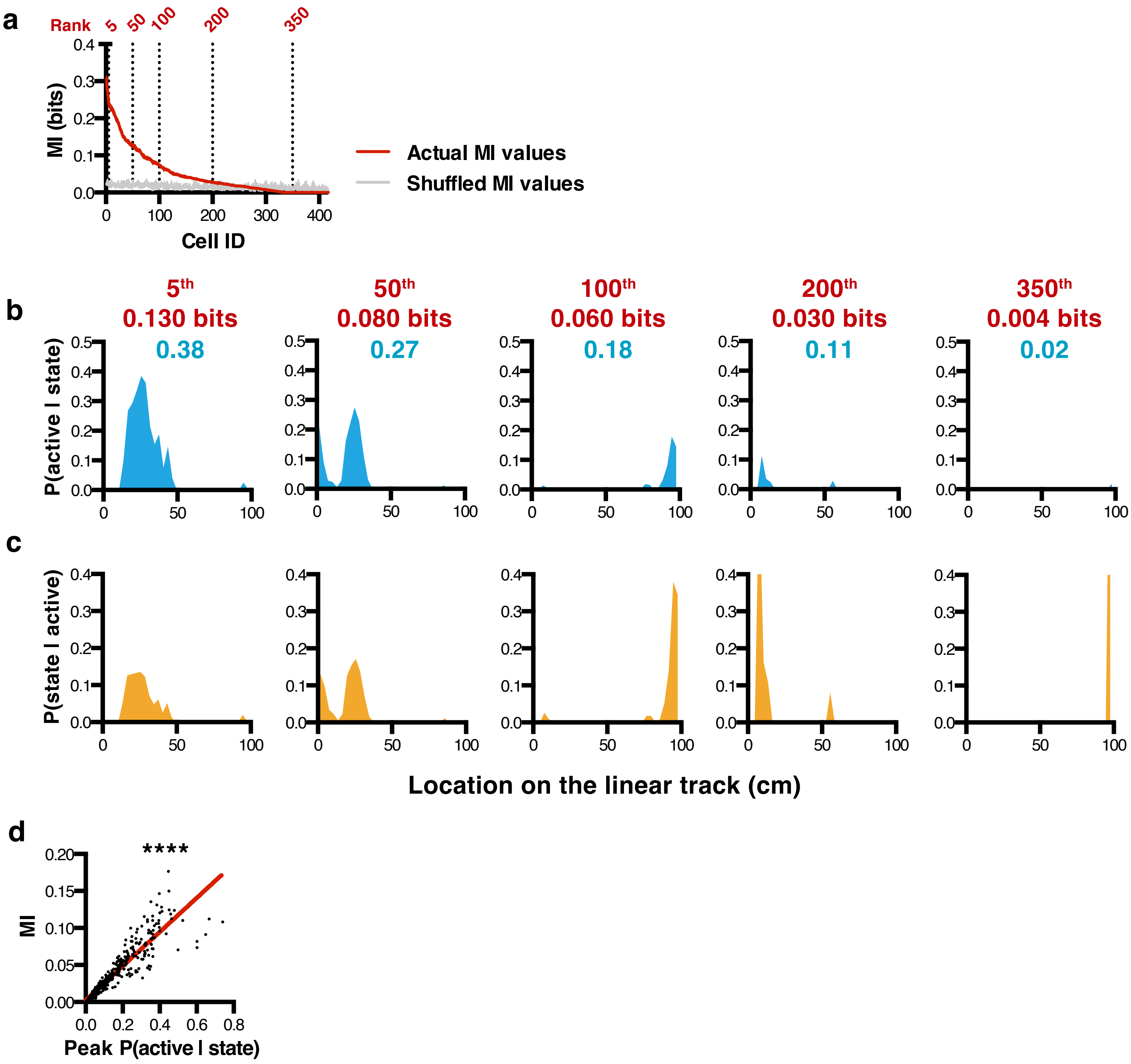


**Supplementary figure 1. Example place cells in the linear track.** a, MI index for every neuron (red), sorted from highest to lowest value, and corresponding cell rank (top) as well as MI value obtained from a shuffled calcium trace (light gray). Thickness of curves represents 95% confidence interval. Example tuning curves of likelihood (b) and posterior (c) probability density functions. Corresponding MI (red) and peak P(active | state) values (blue) are indicated on the top of each panel. d, relationship between peak joint probability and MI index.

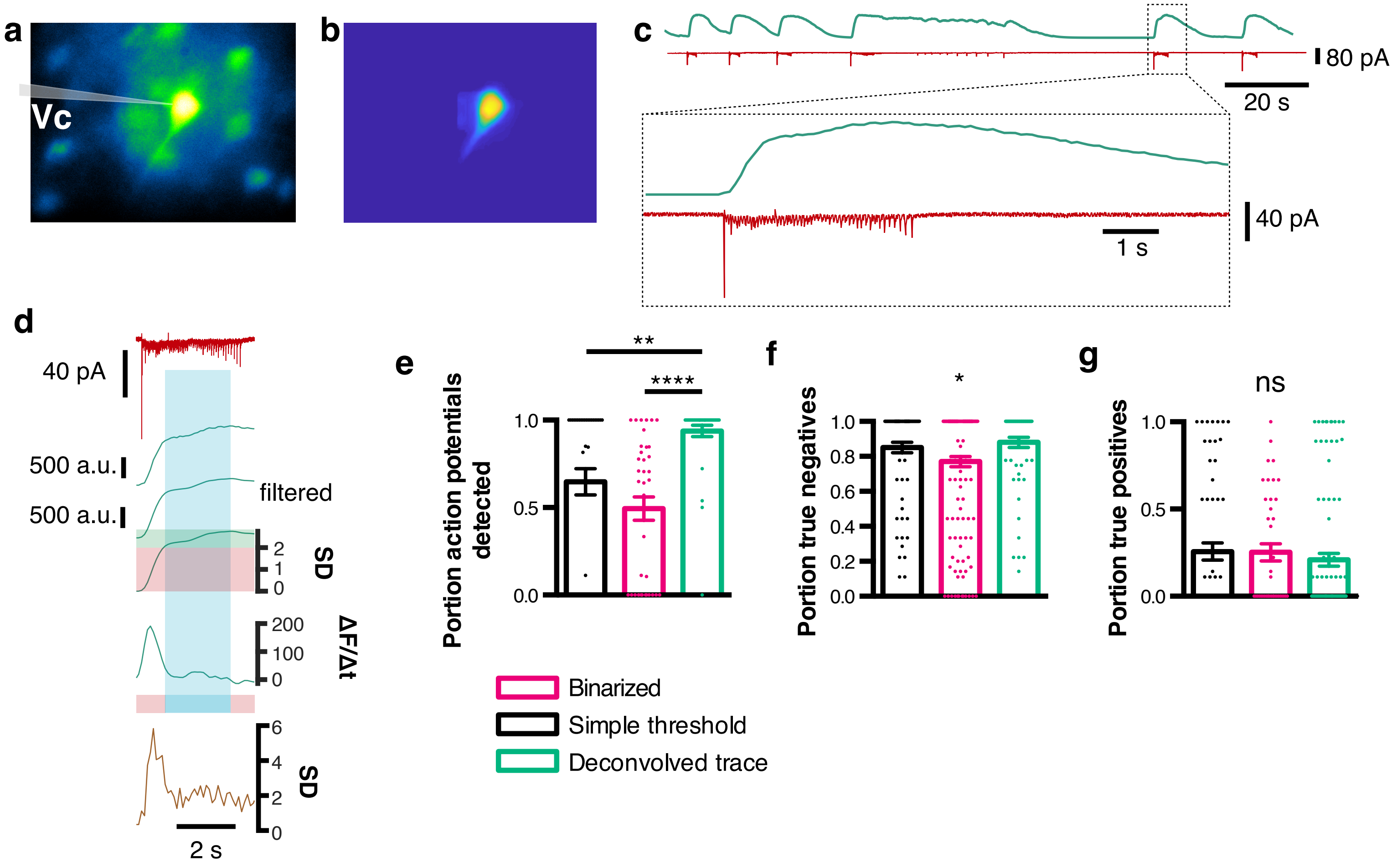


**Supplementary figure 2. Accuracy of binarizing methods in vitro.** a, maximum projection of calcium activity from a dorsal hippocampus section containing pyramidal cell neurons. Patch pipette overlaid to indicate which neuron was recorded. b, spatial footprint of the recorded neuron. c, calcium activity (green) and corresponding membrane currents in cell-attached configuration. d, example currents during a burst of activity (red trace, top) and corresponding calcium traces (green traces) used during the binarizing process, as well as corresponding deconvolution signal (ochre trace, bottom). e, portion of action potentials successfully detected. f, portion of true negatives (epochs labeled as inactive while there was indeed no action potential). g, portion of true positives (epochs alabeled as active while there was indeed at least one action potential).

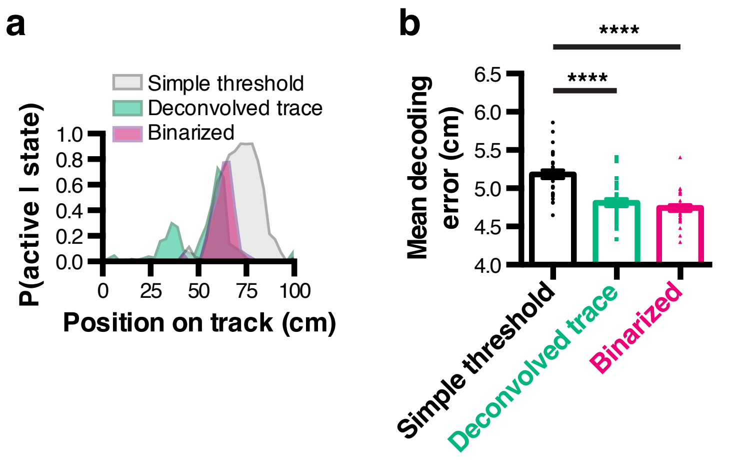


**Supplementary figure 3. Effect of binarizing method on tuning curves and decoding accuracy.** a, tuning curves computed using different binarizing methods. b, mean decoding error derived from different binarizing methods
